# The Mutational Signature Comprehensive Analysis Toolkit (musicatk) for the discovery, prediction, and exploration of mutational signatures

**DOI:** 10.1101/2020.11.17.385864

**Authors:** Aaron Chevalier, Shiyi Yang, Zainab Khurshid, Nathan Sahelijo, Tong Tong, Jonathan Huggins, Masanao Yajima, Joshua D. Campbell

## Abstract

Mutational signatures are patterns of somatic alterations in the genome caused by carcinogenic exposures or aberrant cellular processes. To provide a comprehensive workflow for preprocessing, analysis, and visualization of mutational signatures we created the Mutational Signature Comprehensive Analysis Toolkit (*musicatk*) package. *musicatk* enables users to select different schemas for counting mutation types and easily combine count tables from different schemas. Multiple distinct methods are available to deconvolute signatures and exposures or to predict exposures in individual samples given a pre-existing set of signatures. Additional exploratory features include the ability to compare signatures to the COSMIC database, embed tumors in two dimensions with UMAP, cluster tumors into subgroups based on exposure frequencies, identify differentially active exposures between tumor subgroups and plot exposure distributions across user-defined annotations such as tumor type. Overall, *musicatk* will enable users to gain novel insights into the patterns of mutational signature observed in cancer cohorts.

## Introduction

Somatic mutations to the genome can be caused by exposure to environmental carcinogens or aberrant cellular processes^1,2^. A “mutational signature” is a specific pattern of mutation types caused by a particular mutational process. The set of mutations observed in a single tumor genome can be the result of multiple mutational processes active during the course of tumor development. Therefore, deconvolution is needed to determine which signatures are present across a group of tumor genomes as well as the level of each signature in each individual tumor. Recently, the Pan-Cancer Analysis of Whole Genomes (PCAWG) Consortium characterized a large cohort of whole-exome and whole-genome samples with single-base-substitution (SBS), doublet-base-substitution (DBS), and small insertion-and-deletion (INDEL) mutational schemas using NMF-based methods^3^. While some software packages have been previously developed to perform mutational signature inference, they cannot quantify the latest set of mutation schema from COSMIC^3^. These packages also lack functionality for comprehensive exploratory analysis or have limited functionality for predicting exposures to pre-defined signatures in new samples^3–9^. The *musicatk* package provides functionality to streamline the steps of mutational inference and has several additional features to enhance exploratory analysis beyond what is available in other packages **(Supplementary Table 1)**. We provide an overview of this functionality and present an exploratory analysis of tumors from The Cancer Genome Atlas (TCGA).

## Methods

### Importing and processing of mutations

The major steps of mutational signature inference are 1) importing the variants, 2) building and combining count tables based on different types of mutation schema, 3) performing discovery and/or prediction of signatures and exposures, and 4) using visualization for exploratory analysis of the results **(Figure 1)**. For the first step, the *musicatk* package has functions to read mutations from various input formats. Mutation Annotation Formats (MAFs) are read from files or from the R object *MAF* created by the *maftools* package. Variant Call Formats (VCFs) can be read from files or R classes defined in the *VariantAnnotation* package. Additionally, variant information stored in a *data*.*frame* or *data*.*table* can also be used as input. To streamline the processing of variants, mutation profiles from multiple tumors in different formats can be automatically read and combined into the *musica* object **(Supplementary Figure 1)**.

**Figure 1.**
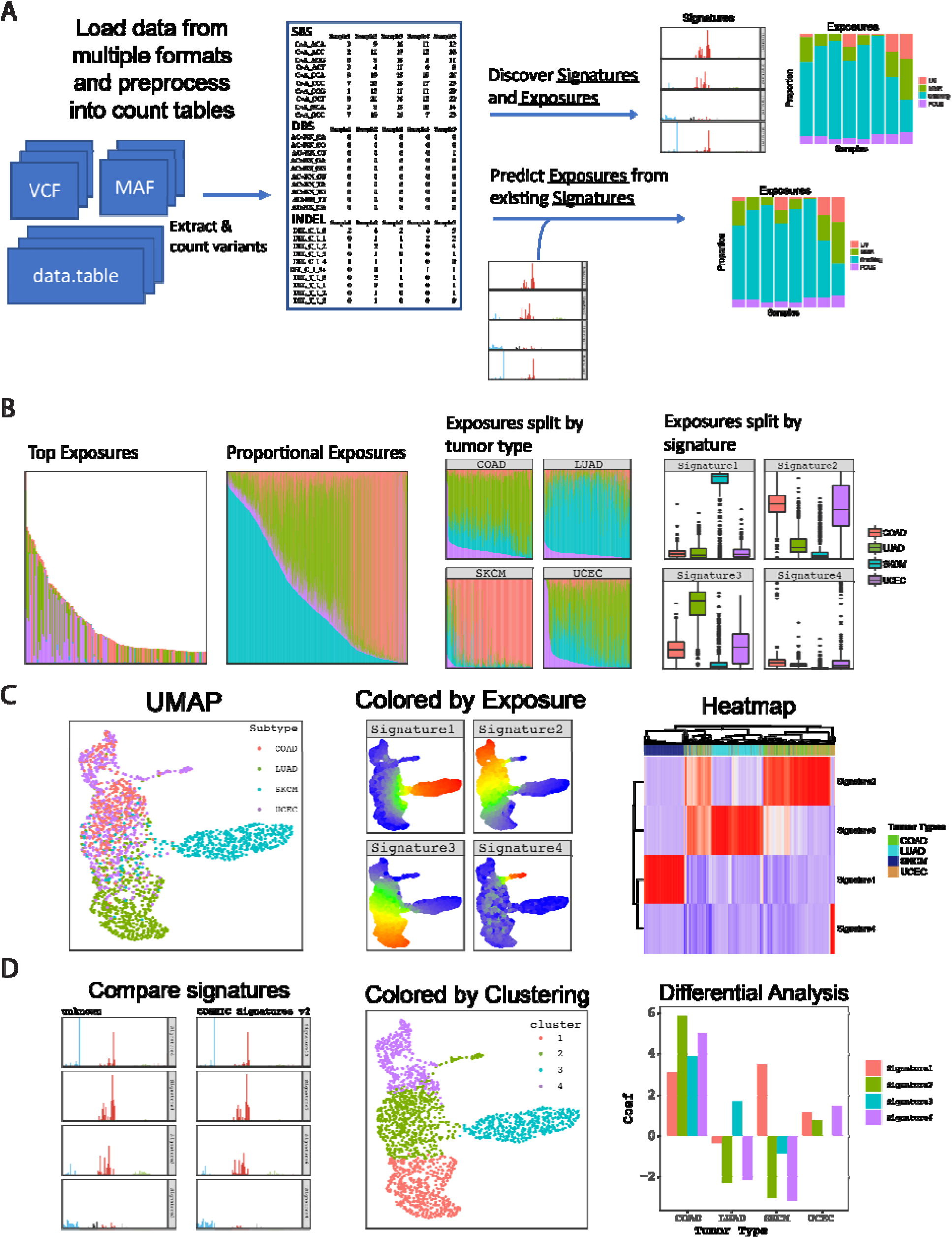
Overview of workflow for mutational signature discovery/prediction, visualization and analysis. **A)** Workflow allows for loading and combining data from multiple sources, de-novo discovery of signature and exposures, and prediction of exposures from existing signatures. **B)** The same sample exposures are plotted (subset to top samples), proportional exposures (signature exposures sum to 1), split up by tumor type, and split up by signature. **C)** Additional visualizations include the ability to plot samples in two dimensions using UMAP. In the UMAP, tumors can be colored by annotations such as tumor type or levels of exposure for each signature. Heatmaps can be used to show the relatively levels of signature exposures in each sample along with sample annotations. **D)** Downstream analysis tools include automated comparison to COSMIC signatures, clustering of tumors by exposure with K-means, and differential analysis of exposures between groups of tumors.

We also include functions that automatically parse different types of mutation motifs from each tumor genome and create count tables that are used in downstream analysis. Tables that can be calculated by *musicatk* include SBS into 96 motifs, SBS with transcription strand orientation into 192 motifs, SBS with replication strand orientation into 192 motifs, DBS into 78 motifs, and INDELs into 83 motifs. Custom mutation count tables can also be defined by the user and added to the object. Importantly, multiple tables can be concatenated to create composite mutation schema tables. For example, users can combine the SBS-96 motif, DBS-78 motif, and INDEL-83 tables and perform downstream analyses, similar to process that was used to create the PCAWG Composite signatures^3^.

### Discovery of mutational signatures

Deconvolution is the process of decomposing a matrix of mutation counts per tumor into a matrix of signatures and exposures. The *Signature* matrix contains the probability of each mutation motif in each signature and the *Exposure* matrix contains the estimated level of each exposure in each tumor sample. *musicatk* supports both Latent Dirichlet Allocation (LDA)^10^ from the *topicmodels* package and Non-Negative Matrix Factorization (NMF)^11^ from the *NMF* package. Both algorithms can be applied to any count table. One challenging aspect of mutational signature discovery is determining the appropriate number of signatures (i.e., the value of *K*). To facilitate the comparison of models with different choices of *K, musicatk* provides a wrapper that allows users to apply deconvolution algorithms with different values of *K* and then compare the results with metrics such as reconstruction error (NMF and LDA), log-likelihood (LDA), or perplexity (LDA).

### Prediction of mutational signatures

Prediction of exposures for existing signatures can be performed on any count table given that the mutation motif schema is the same. We include wrappers for tools such as *deconstructSigs* and *decompTumor2Sig*^4,8^. We also implement a Bayesian algorithm based on LDA where exposures are estimated using a fixed set of signatures **(Supplementary File 1)**. To allow for prediction using previously defined signatures from the COSMIC database^3^, we include objects for COSMIC V2 signatures (SBS-96 motif schema) and COSMIC V3 signatures (SBS-96, DBS-78, and INDEL-83 motif schemas). One challenge in the prediction of existing signatures in new tumors is that not all signatures will be present in the new dataset. Including non-active signatures may cause additional noise in the estimates for signatures that are present within the dataset. Since signatures that are present in moderate levels across many tumors or highly present in a small number of tumors are more likely to be active in the dataset^3^, we implemented a two-step procedure to choose the subset of active signatures within a dataset. In the first step, exposures are estimated using all signatures and active signatures are chosen if they pass a threshold in a minimum number of samples (e.g. have an exposure of at least 0.1 in at least 30% of samples or an exposure of 0.7 in at least 2 samples). In the second-step, signatures are estimated using only the active signatures. Users can perform the 2-step prediction within subgroups of tumors supplying a categorical annotation such as tumor type.

### Visualization

Visualization of mutational signatures and tumor exposures is an important for characterization of mutational processes active in a cohort of tumors. Barplot functions can be used to show the probability of each motifs in each signature and the composition of exposures in each sample. Exposure barplots can be sorted by overall mutation count, one or more exposures, or sample name **(Figure 1B)**. Exposure barplots can be subsetted to show the samples with the highest total mutation counts or exposure levels. Exposures barplots can further be grouped by a sample annotation such as tumor type or by signature. The distributions of exposures can be displayed with box and/or violin plots and grouped by sample annotations. To view relationships between tumors in two dimensions, the Uniform Manifold Approximation and Projections (UMAPs)^12^ algorithm can be used with normalized signature exposures **(Figure 1C)**. The UMAP can be colored by annotations (e.g., tumor type) or the levels of each exposure. Finally, exposures can be viewed in a heatmap along with sample annotations and arranged by hierarchical clustering.

### Down-stream analyses

Functionality is provided for correlating sets of signatures between different signature result objects. This can be used to easily perform comparisons to COSMIC V2 and V3 signatures **(Figure 1D)**. musicatk provides clustering using multiple distance metrics (Cosine, Euclidean, Manhattan, Jaccard, Canberra) in order to group samples based on exposure. These can be plotted on top of a UMAP **(Figure 1D)**. The *ComplexHeatmap* package is used to plot heatmaps showing the relative levels of exposures in samples along with annotations **(Figure 1C)**. K-means can be applied to exposures to cluster tumors into groups. Metrics such as silhouette width and total within some of squares (wss) generated from the *factoextra* package can be used to identify the optimal number of clusters. Differential analysis can be used to identify exposures that are significantly higher or lower between groups of tumors. Differential methods include Wilcoxon rank-sum test for 2-group comparisons as well as Kruskal-Wallis and Negative Binomial Generalized Linear Models (GLMs) for multi-group comparisons **(Figure 1D)**.

## Results

### Single-base substitutions (SBS)

We applied the LDA-based prediction method to predict COSMIC v3 SBS signatures in a Pan-Cancer dataset from TCGA. 39 of the 65 signatures were found to be active in at least one tumor type. A UMAP plot was generated to explore the patterns of signatures across tumors **(Figure 2A)**. Some signatures were present in nearly half of samples, some in a few tumor types, some in single tumor types, and some in subsets of multiple tumor types **(Figure 2, Supplementary Figure 2**). For example, APOBEC signatures (SBS2 and SBS13) were present in a subset of tumors BRCA, CESC, BLCA, and HNSC, and distantly in a subset of LUAD and LUSC tumors. In general, both APOBEC signatures were present in the same samples. The only exceptions were disjoint subsets of BRCA samples with either SBS2 or SBS13. UCEC samples are split into three groups. The bottom samples are clustered separately because of their exposure to the SBS39 signature (unknown origin). The small top cluster includes samples from a few other tumor types including COAD and is defined by high levels of the POLE signature (SBS10 a/b). The left cluster has high levels of a defective DNA mismatch repair signature (SBS44). Two other MMR signatures, SBS15 and SBS21 could distinguish subsets of COAD and STAD tumors which also had higher levels of a third MMR signature, SBS6.

**Figure 2.**
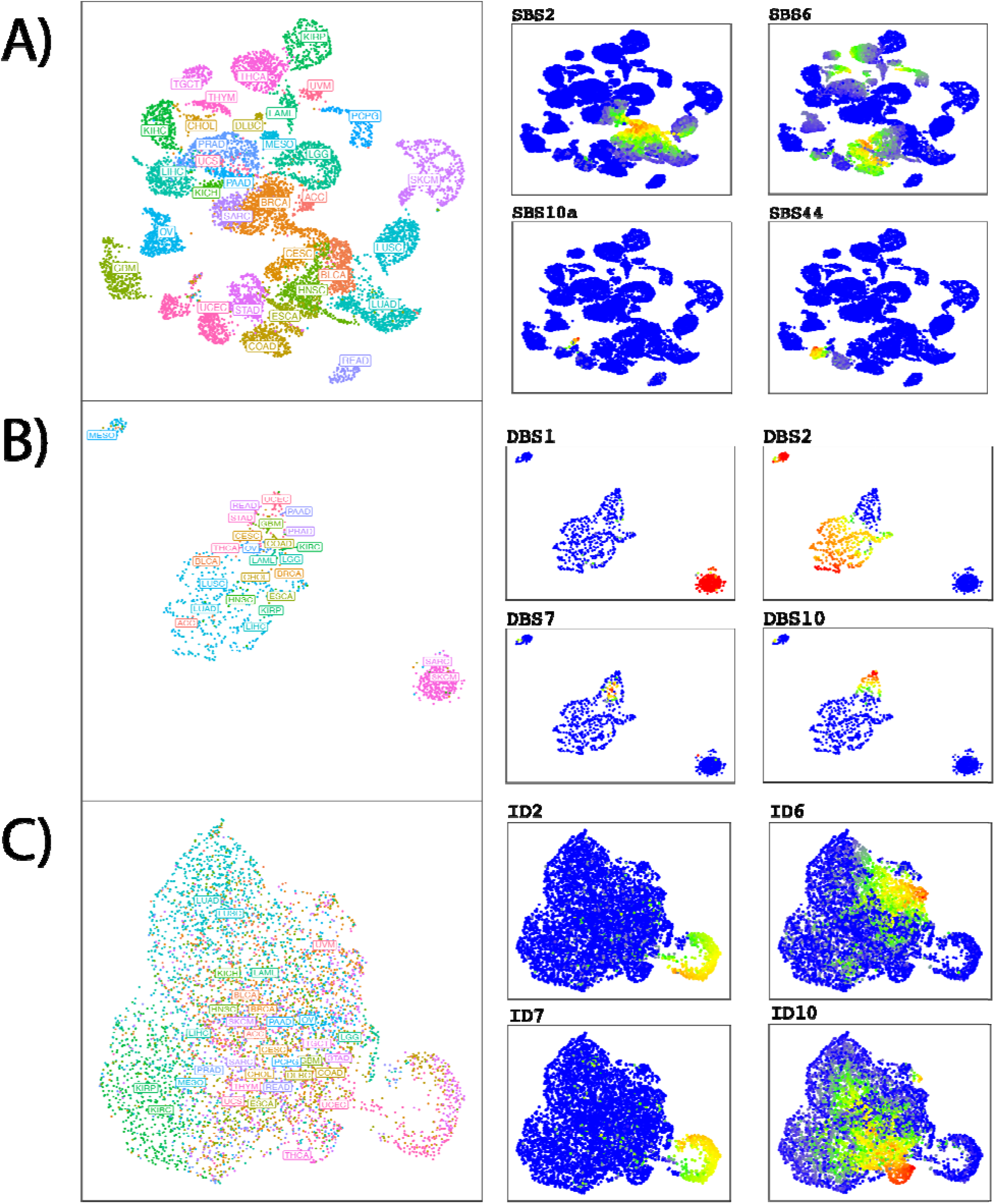
UMAPs of TCGA sample exposure to COSMIC v3 signatures. A UMAP was generated with the *uwot* package usin the exposures predicted from LDA for different mutation types. The left plot is colored by the tumor type while the right plot is colored by exposure levels of each signature (red being high and blue being low). **A)** Examples of Single Base Substitution (SBS) exposures include SBS2, SBS6, SBS10a, SBS44 representing APOBEC, MMR, POLE, and MMR, respectively. **B)** Examples of Double Base Substitution (DBS) exposures include DBS1, DBS2, DBS7, DBS10 representing UV, tobacco, MMR, and MMR, respectively. **C)** Examples of small Insertion and Deletion (INDEL) exposures include ID2, ID6, ID7, ID10 representing spillage during DNA replication, homologous recombination-based DNA damage repair, MMR, and unknown etiology, respectively.

### Double-base substitutions (DBS)

11 of 11 COSMIC v3 DBS signatures were active in TCGA samples. Examining the DBS UMAP showed that DBS2 (tobacco smoke) is active in two of the 3 major clusters, representing ACC, LUAD, LUSC, HNSC, KIRP, LIHC, BLCA, ESCA, and MESO **(Figure 2B, Supplementary Figure 3)**. DBS1 (UV light exposure) is found only in the SARC/SKCM cluster. DBS10 (defective DNA mismatch repair) is predominantly found in tumors from READ, PAAD, UCEC, and STAD and active in different sets of tumors from DBS1 and DBS2. DBS7 is also caused by defective DNA mismatch repair and mostly active in different sets of tumors from DBS10. The remaining signatures are present in mixed subsets of tumor types.

### Insertions and deletions (INDELs)

All 17 COSMIC v3 INDEL signatures were predicted to be active in TCGA samples **(Figure 2C, Supplementary Figure 4)**. ID3 (tobacco smoking) was predominantly active in tumors from LUAD and LUSC. ID6 (defective DNA repair) was highly active in a distinct subset of samples containing mix of tumor types such as BRCA, OV, and STAD. High levels of ID10 (unknown etiology) defined a unique group of samples that were enriched for tumors from THCA and SARC. A distinct group of mixed tumor types was defined by different levels of activity for ID2 (defective DNA replication) and ID7 (defective DNA mismatch repair) suggesting that these aberrant processes may often co-occur.

In conclusion, the *musicatk* package provides a comprehensive set of preprocessing utilities, access to several discovery and prediction tools, and functions for downstream analysis of the patterns of mutational signatures in a cohort of tumors.

## Acknowledgements

This work was funded by the Informatics Technology for Cancer Research (ITCR) R21 CA226188-01A1 (J.D.C., M.Y.) and by the National Institute of General Medical Sciences of the National Institutes of Health T32GM100842 (Z.K., N.S., T.S.).

---

**Supplementary Table 1.**
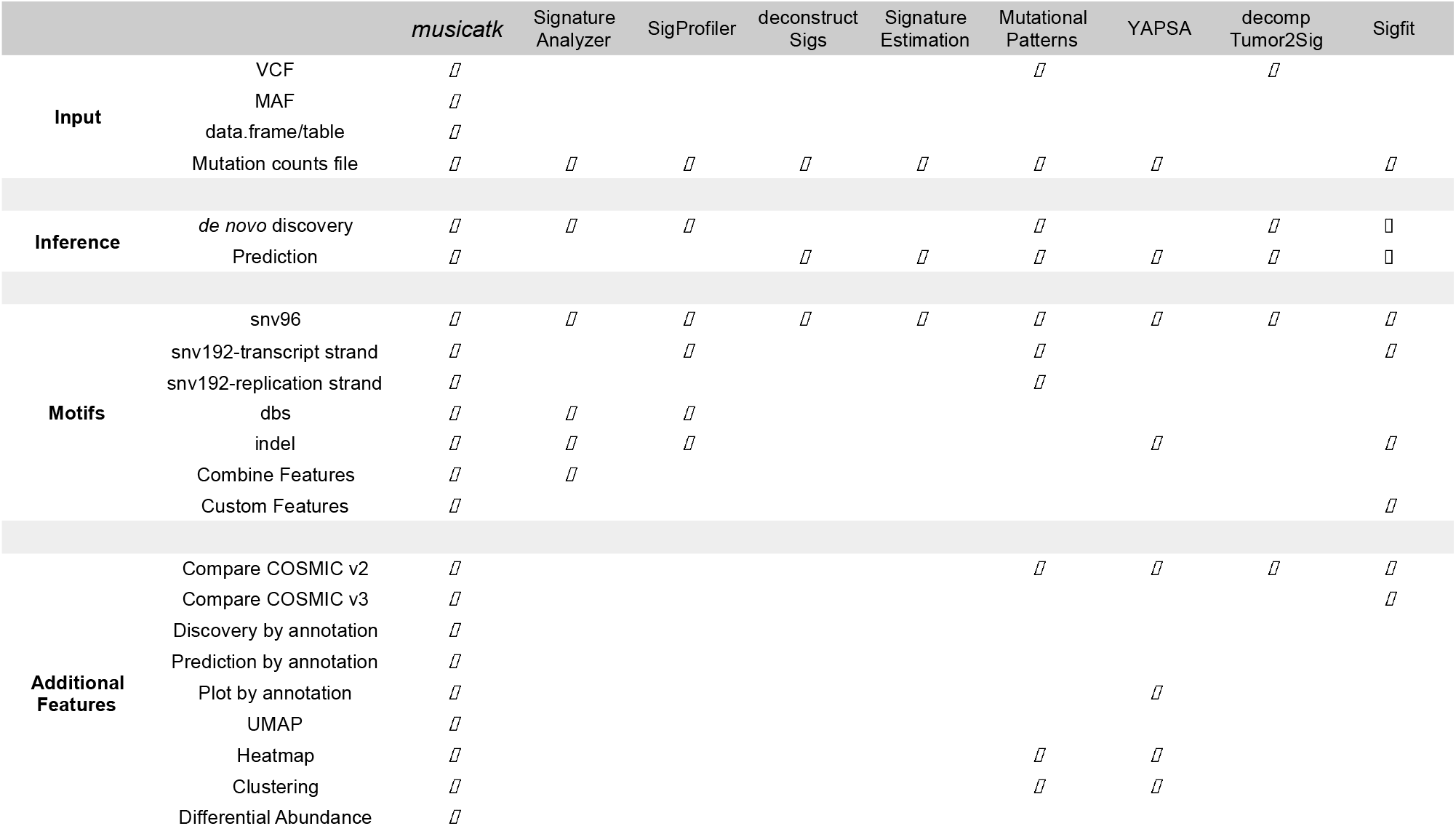
Comparison of features across mutational signature packages.

**Supplementary Figure 1.**
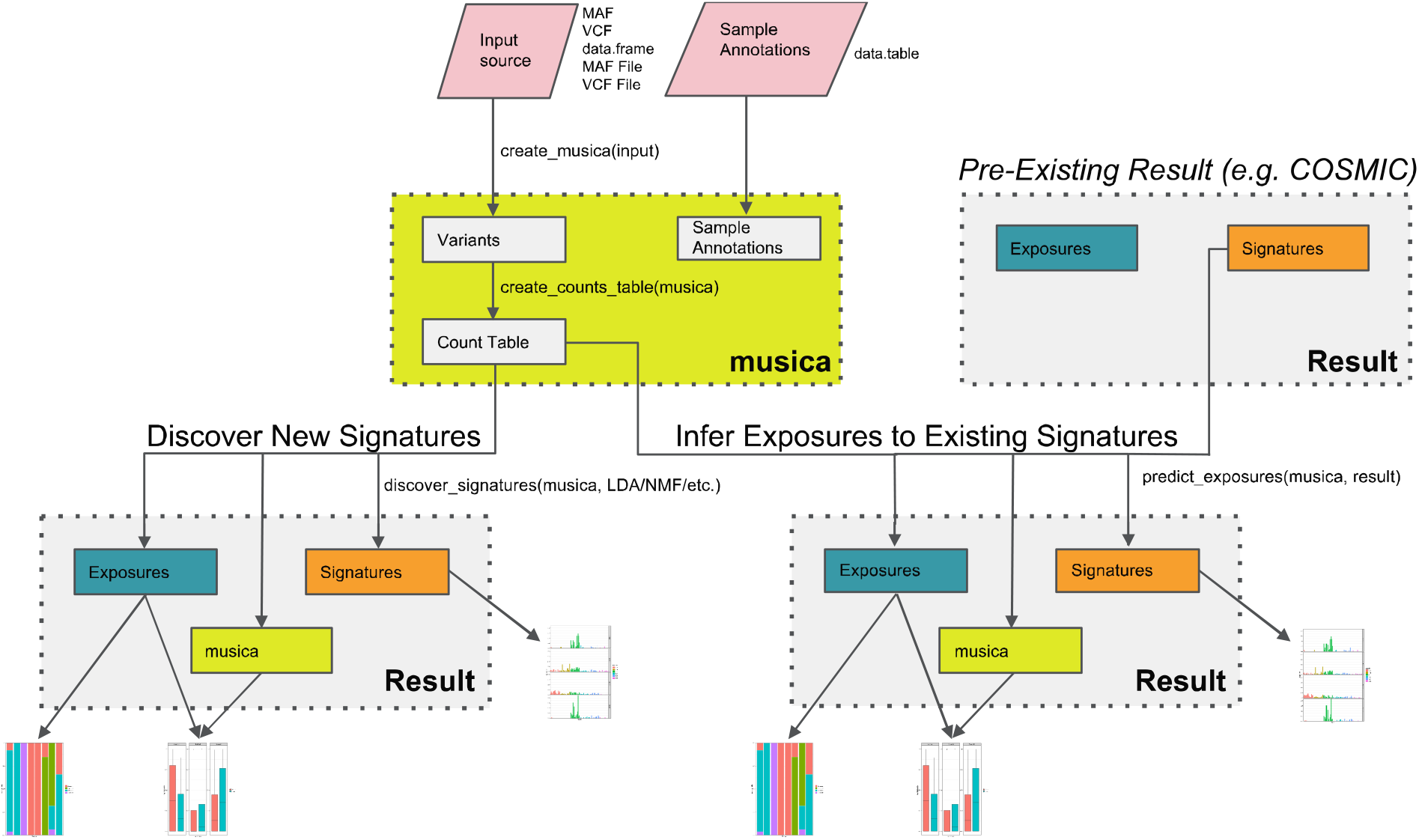
Overview of the class structure in the *musicatk* package. This *musica* class is used as the main object for building count tables (SBS96, DBS78, IND83, SBS192, etc.) and for storing count tables for all variant classes. A *musica* object is used as input into the discovery or prediction functions. A *mustica_result* object is output from discover and prediction functions and used to store the variant tables along with the estimated signatures and exposures matrices. Existing signatures contained in a *mustica_result* object can be used to predict exposures in a new dataset. Result objects are available for COSMIC v2 and COSMIC v3 signatures in the package. User-generated *mustica_result* objects are used as input into all down-stream plotting and analysis functions.

**Supplementary Figure 2.**
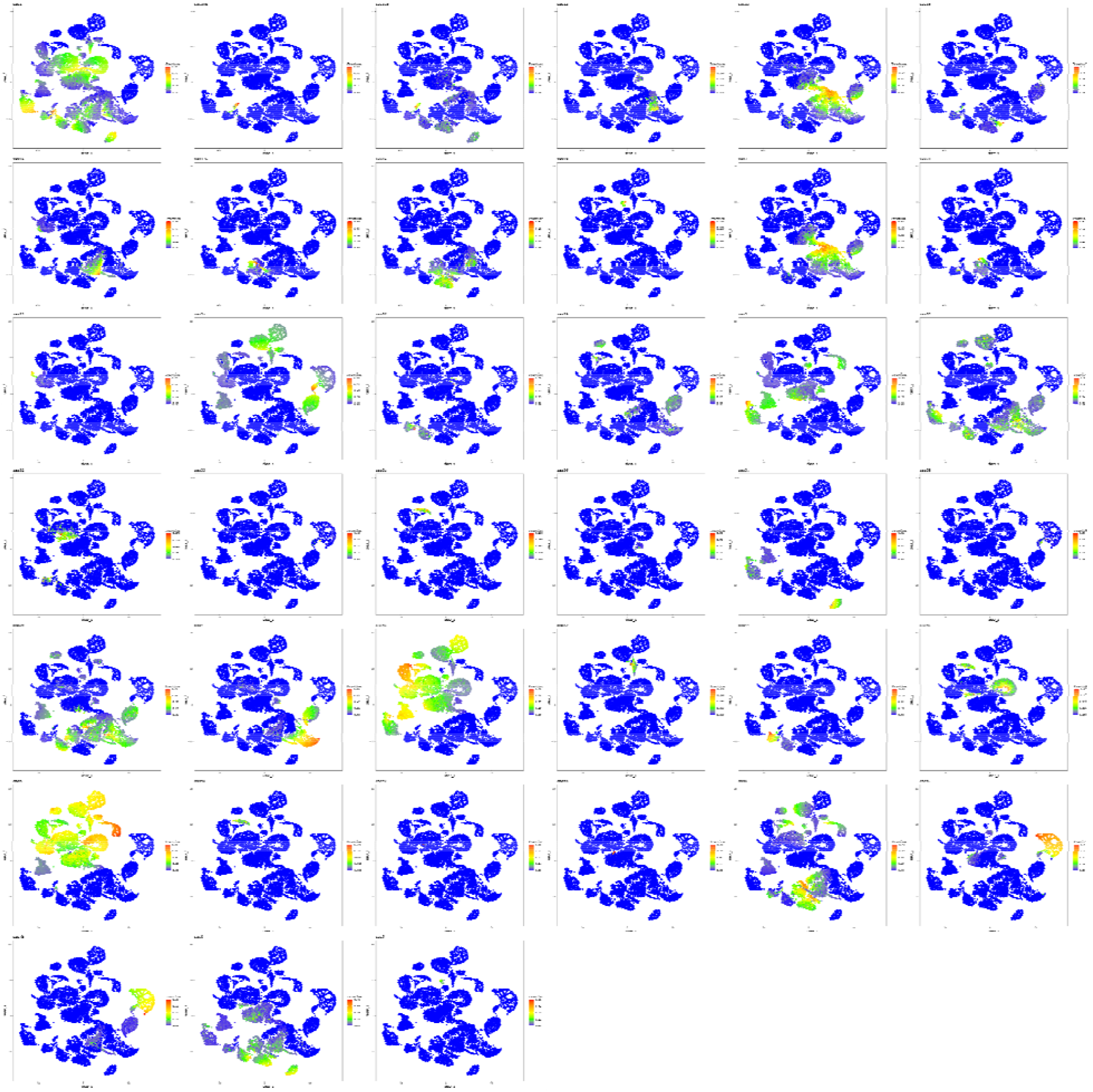
UMAP colored by the predicted exposures from COSMIC v3 SBS signature.

**Supplementary Figure 3.**
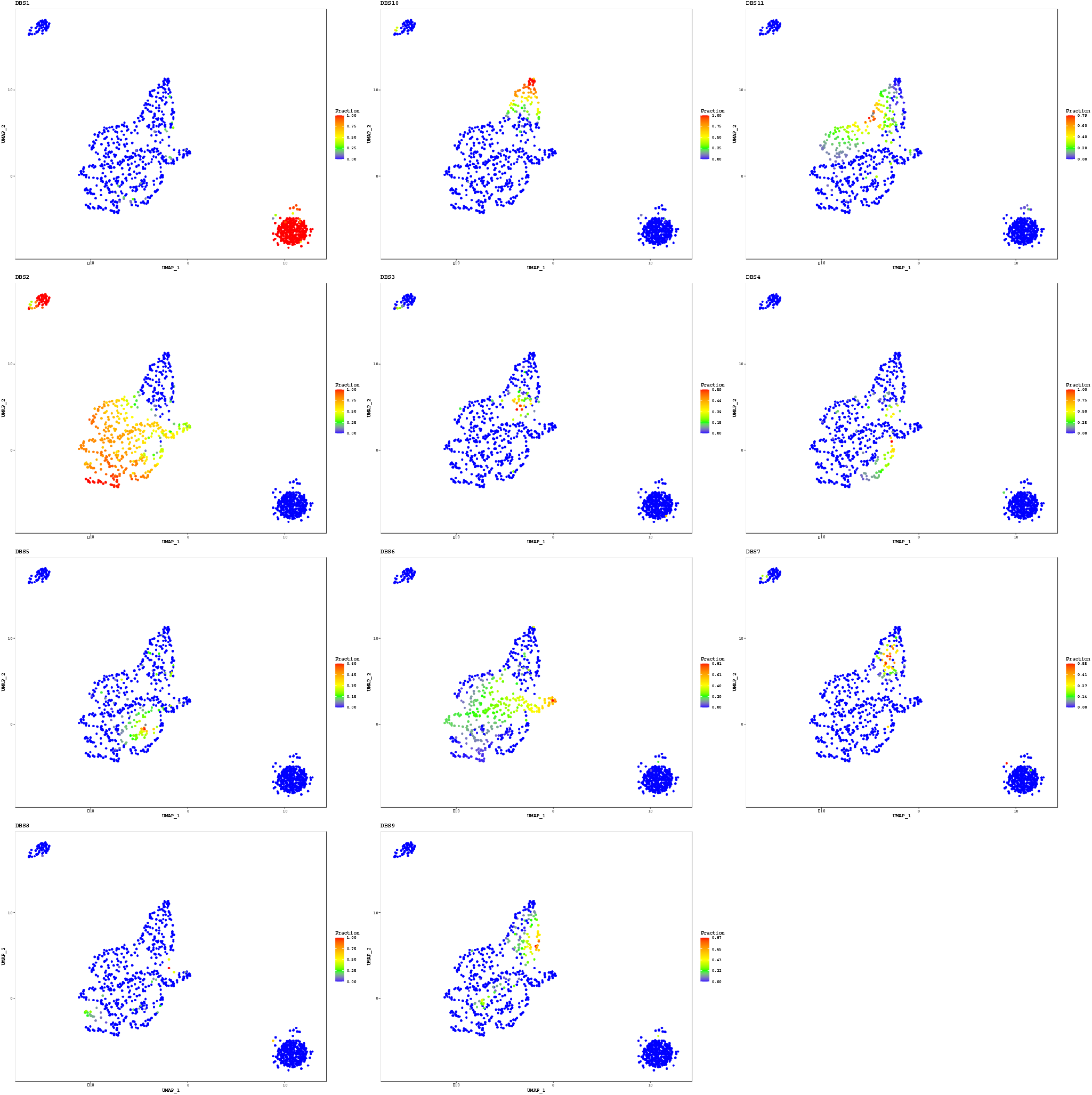
UMAP colored by the predicted exposures from COSMIC v3 DBS signatures.

**Supplementary Figure 4.**
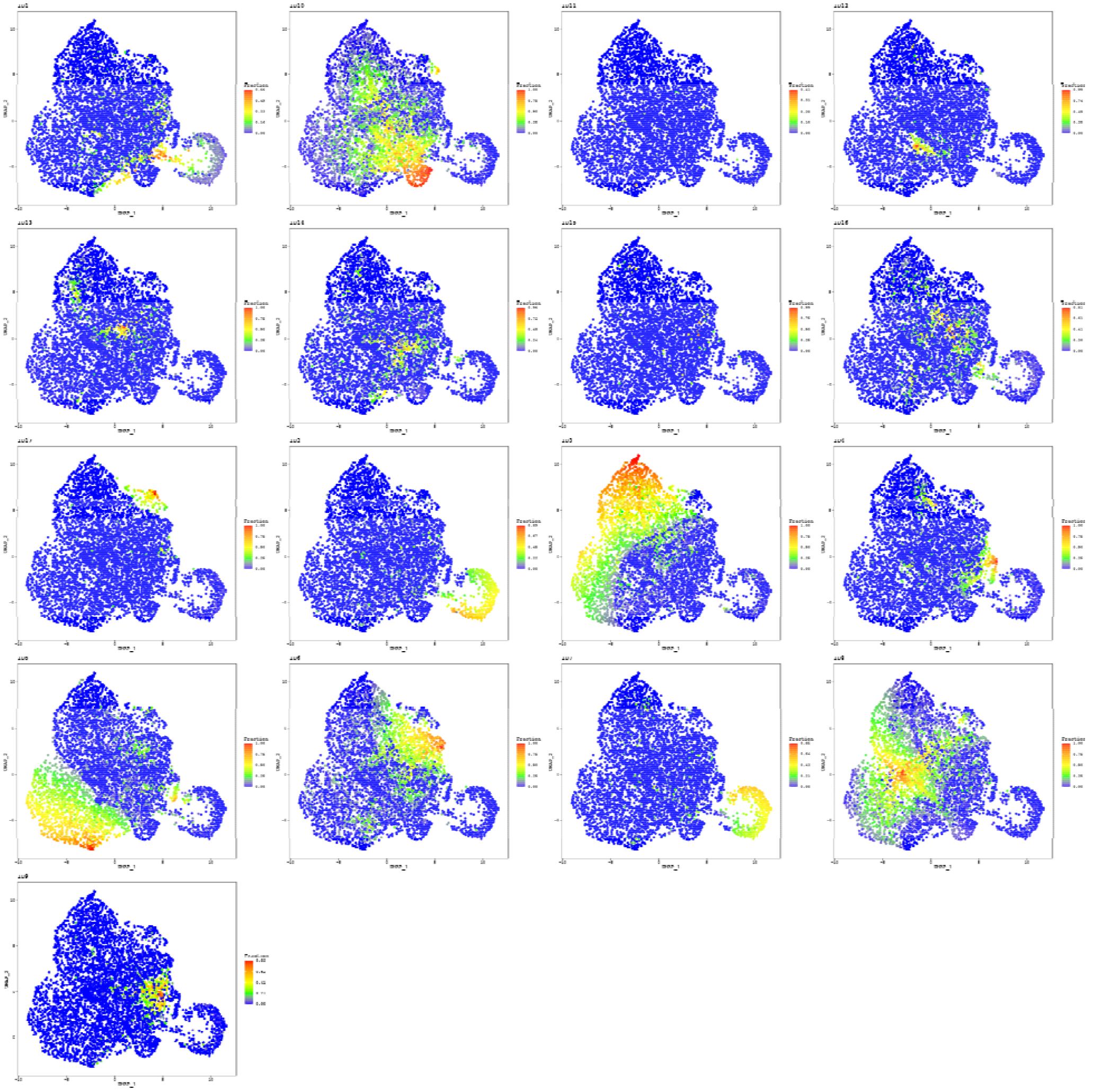
UMAP colored by the predicted exposures from COSMIC v3 INDEL signatures.

## References

1. Nik-Zainal, S. et al. Landscape of somatic mutations in 560 breast cancer whole-genome sequences. Nature 534, 47–54 (2016).

2. Alexandrov, L. B. et al. Signatures of mutational processes in human cancer. Nature 500, 415 (2013).

3. Alexandrov, L. B. et al. The repertoire of mutational signatures in human cancer. Nature 578, 94–101 (2020).

4. Rosenthal, R., McGranahan, N., Herrero, J., Taylor, B. S. & Swanton, C. deconstructSigs: delineating mutational processes in single tumors distinguishes DNA repair deficiencies and patterns of carcinoma evolution. Genome Biol. 17, 31 (2016).

5. Teresa Przytycka Research Page. Available at: https://www.ncbi.nlm.nih.gov/CBBresearch/Przytycka/index.cgi#signatureestimation. (Accessed: 16th November 2020)

6. Blokzijl, F., Janssen, R., van Boxtel, R. & Cuppen, E. MutationalPatterns: Comprehensive genome-wide analysis of mutational processes. Genome Med. 10, 33 (2018).

7. YAPSA: Yet Another Package for Signature Analysis version 1.16.0 from Bioconductor. Available at: https://rdrr.io/bioc/YAPSA/. (Accessed: 16th November 2020)

8. Krüger, S. & Piro, R. M. DecompTumor2Sig: Identification of mutational signatures active in individual tumors. BMC Bioinformatics 20, 152 (2019).

9. Gori, K. & Baez-Ortega, A. sigfit: flexible Bayesian inference of mutational signatures. bioRxiv 372896 (2018). doi:10.1101/372896

10. Blei, D. M., Ng, A. Y. & Jordan, M. I. Latent Dirichlet Allocation. J. Mach. Learn. Res. 3, 993–1022 (2003).

11. Lee, D. D. & Seung, H. S. Learning the parts of objects by non-negative matrix factorization. Nature 401, 788–791 (1999).

12. McInnes, L., Healy, J. & Melville, J. UMAP: Uniform Manifold Approximation and Projection for Dimension Reduction. arXiv (2018).

