## Supplementary File 1 for "The Mutational Signature Comprehensive Analysis Toolkit (musicatk) for the discovery, prediction, and exploration of mutational signatures"

### 1 Generative process

1. For each sample $s$, $s \in \{1, 2, ..., S\}$:
   (a) Generate signature proportions $\theta_s \sim Dir^K(\omega)$
   (b) For each mutation motif $x_n$, $n \in \{1, 2, ..., N_s\}$:
      i. Generate signature type $z_n \sim Multinomial(\theta_s)$
      ii. Generate mutation type $x_n \sim Multinomial(\eta_{z_n})$

$S$: Number of samples

$N_s$: Number of mutations in sample $s$

$K$: Number of signatures

$\eta_k$: mutation distribution for signature $k$, known *apriori*

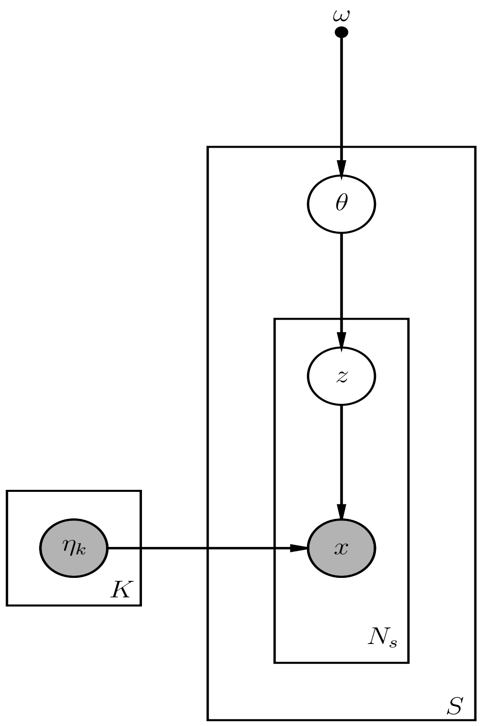


**Plate diagram**

### 2 Likelihood

$$P(x, \boldsymbol{\theta}, \boldsymbol{z}|\omega, \boldsymbol{\eta}) = \prod_{s=1}^{S} P_{Dir}(\theta_s|\omega) \left[ \prod_{n=1}^{N_s} P_{Mult}(z_n|\theta_s) P_{Mult}(x_n|\eta_{z_n}) \right] \quad (1)$$

, where $\omega$ is the hyperparameter for $\theta$

#### 3 Inference

Log-likelihood is:

$$\log P(x, \boldsymbol{\theta}, \boldsymbol{z}|\omega, \boldsymbol{\eta}) = \sum_{s=1}^{S}\left[\log P_{Dir}(\theta_s|\omega) + \left(\sum_{n=1}^{N_s}\log P_{Mult}(z_n|\theta_s) + \log P_{Mult}(x_n|\eta_{z_n})\right)\right] \tag{2}$$

Log-likelihood for mutations within one sample can be written as:

$$\log P = \log P(\theta_s|\omega) + \left(\sum_{n=1}^{N_s}\log P(z_n|\theta_s) + \log P(x_n|\eta_{z_n})\right) \tag{3}$$

We use variational inference to estimate parameters, and use variational distributions $q(\theta, \boldsymbol{z}|\alpha, \boldsymbol{\pi})$ to approximate and decouple $\theta$ and $\boldsymbol{z}$. The variational distribution is specified as following:

$$q(\boldsymbol{\theta_s}, \boldsymbol{z}|\alpha_s, \boldsymbol{\pi}) = \prod_{s=1}^{S} q_{Dir}(\theta_s|\alpha_s)(\prod_{n=1}^{N_s} q_{Mult}(z_n|\pi_n)) \tag{4}$$

Using Jensen's inequality, we have:

$$\begin{aligned}
\log P(x|\omega, \boldsymbol{\eta}) &= \log \int_{\boldsymbol{\theta}}\int_{\boldsymbol{z}} P(x, \boldsymbol{\theta}, \boldsymbol{z}|\omega, \boldsymbol{\eta}) d\boldsymbol{z} d\boldsymbol{\theta} \\
&= \log \int_{\boldsymbol{\theta}}\int_{\boldsymbol{z}} \frac{P(x, \boldsymbol{\theta}, \boldsymbol{z}|\omega, \boldsymbol{\eta})}{q(\boldsymbol{\theta}, \boldsymbol{z}|\boldsymbol{\alpha}, \boldsymbol{\pi})} q(\boldsymbol{\theta}, \boldsymbol{z}|\boldsymbol{\alpha}, \boldsymbol{\pi}) d\boldsymbol{z} d\boldsymbol{\theta} \\
&\geq \int_{\boldsymbol{\theta}}\int_{\boldsymbol{z}} \log \frac{P(x, \boldsymbol{\theta}, \boldsymbol{z}|\omega, \boldsymbol{\eta})}{q(\boldsymbol{\theta}, \boldsymbol{z}|\boldsymbol{\alpha}, \boldsymbol{\pi})} q(\boldsymbol{\theta}, \boldsymbol{z}|\boldsymbol{\alpha}, \boldsymbol{\pi}) d\boldsymbol{z} d\boldsymbol{\theta} \\
&= \int_{\boldsymbol{\theta}}\int_{\boldsymbol{z}} \left[\log P(x, \boldsymbol{\theta}, \boldsymbol{z}|\omega, \boldsymbol{\eta}) - \log q(\boldsymbol{\theta}, \boldsymbol{z}|\boldsymbol{\alpha}, \boldsymbol{\pi})\right] q(\boldsymbol{\theta}, \boldsymbol{z}|\boldsymbol{\alpha}, \boldsymbol{\pi}) d\boldsymbol{z} d\boldsymbol{\theta} \\
&= \int_{\boldsymbol{\theta}}\int_{\boldsymbol{z}} \log P(x, \boldsymbol{\theta}, \boldsymbol{z}|\omega, \boldsymbol{\eta}) q(\boldsymbol{\theta}, \boldsymbol{z}|\boldsymbol{\alpha}, \boldsymbol{\pi}) d\boldsymbol{z} d\boldsymbol{\theta} - \int_{\boldsymbol{\theta}}\int_{\boldsymbol{z}} \log q(\boldsymbol{\theta}, \boldsymbol{z}|\boldsymbol{\alpha}, \boldsymbol{\pi}) q(\boldsymbol{\theta}, \boldsymbol{z}|\boldsymbol{\alpha}, \boldsymbol{\pi}) d\boldsymbol{z} d\boldsymbol{\theta} \\
&= E_{q(\boldsymbol{\theta}, \boldsymbol{z}|\boldsymbol{\alpha}, \boldsymbol{\pi})}[\log P(x, \boldsymbol{\theta}, \boldsymbol{z}|\omega, \boldsymbol{\eta})] - E_{q(\boldsymbol{\theta}, \boldsymbol{z}|\boldsymbol{\alpha}, \boldsymbol{\pi})}[\log q(\boldsymbol{\theta}, \boldsymbol{z}|\boldsymbol{\alpha}, \boldsymbol{\pi})]
\end{aligned} \tag{5}$$

The right-hand side of equation(5) is the evidence lower bound which we will refer to as $L(\boldsymbol{\alpha}, \boldsymbol{\pi}; \omega)$. To maximize the lower bound is equivalent to minimizing the KL divergence between the variational posterior distributions and true posterior distributions.

We use $q$ to denote $q(\boldsymbol{\theta}, \boldsymbol{z}|\boldsymbol{\alpha}, \boldsymbol{\pi})$ in subscripts and expand evidence lower bound by plugging in equation(2) and equation(4) as follows:

$$\begin{aligned}
L(\boldsymbol{\alpha}, \boldsymbol{\pi}; \omega) &= E_q[\log P(x, \boldsymbol{\theta}, \boldsymbol{z}|\omega, \boldsymbol{\eta})] - E_q[\log q(\boldsymbol{\theta}, \boldsymbol{z}|\boldsymbol{\alpha}, \boldsymbol{\pi})] \\
&= E_q\left[\sum_{s=1}^{S}\left(\log P(\theta_s|\omega) + \sum_{n=1}^{N_s}(\log P(z_n|\theta_s) + \log P(x_n|\eta_{z_n}))\right)\right] \\
&\quad - E_q\left[\sum_{s=1}^{S}\left(\log q(\theta_s|\alpha_s) + \sum_{n=1}^{N_s}\log q(z_n|\pi_n)\right)\right],
\end{aligned} \tag{6}$$

where $\alpha_s$ is the parameter vector of length $K$ in the Dirichlet distribution for $\theta_s$, $\pi_n$ is the parameter vector of length $K$ in the multinomial distribution for $z_n$.

We can maximize the lower bound with respect to variational parameters within each sample. The lower bound within one sample can then be written and expanded as:

$$
\begin{aligned}
L_s(\boldsymbol{\alpha}, \boldsymbol{\pi}; \omega) &= E_q\left[\log P(\theta_s|\omega) + \sum_{n=1}^{N_s}(\log P(z_n|\theta_s) + \log P(x_n|\eta_{z_n}))\right] \\
&- E_q\left[\log q(\theta_s|\alpha_s) + \sum_{n=1}^{N_s}\log q(z_n|\pi_n)\right] \\
&= E_q\left[\log\Gamma(\sum_{k=1}^{K}\omega_k) - \sum_{k=1}^{K}\log\Gamma(\omega_k) + \sum_{k=1}^{K}(\omega_k - 1)\log\theta_{k,s}\right] \\
&+ E_q\left[\sum_{k=1}^{K}\sum_{n=1}^{N_s} I(z_n = k)\log\theta_{k,s}\right] \\
&+ E_q\left[\sum_{k=1}^{K}\sum_{n=1}^{N_s}\sum_{t=1}^{T} I(z_n = k)x_n^t\log\eta_k^t\right] \\
&- E_q\left[\log\Gamma(\sum_{k=1}^{K}\alpha_{s,k}) - \sum_{k=1}^{K}\log\Gamma(\alpha_{s,k}) + \sum_{k=1}^{K}(\alpha_{s,k} - 1)\log\theta_{s,k}\right] \\
&- E_q\left[\sum_{k=1}^{K}\sum_{n=1}^{N_s} I(z_n = k)\log\pi_{n,k}\right] \\
&= \left[\log\Gamma(\sum_{k=1}^{K}\omega_k) - \sum_{k=1}^{K}\log\Gamma(\omega_k) + \sum_{k=1}^{K}(\omega_k - 1)(\Psi(\alpha_k) - \Psi(\sum_{k''=1}^{K}\alpha_{s,k''}))\right] \\
&+ \left[\sum_{k=1}^{K}\sum_{n=1}^{N_s}\pi_{n,k}(\Psi(\alpha_k) - \Psi(\sum_{k''=1}^{K}\alpha_{s,k''}))\right] \\
&+ \left[\sum_{k=1}^{K}\sum_{n=1}^{N_s}\sum_{t=1}^{T}\pi_{n,k}x_n^t\log\eta_k^t\right] \\
&- \left[\log\Gamma(\sum_{k=1}^{K}\alpha_{s,k}) - \sum_{k=1}^{K}\log\Gamma(\alpha_{s,k}) + \sum_{k=1}^{K}(\alpha_{s,k} - 1)(\Psi(\alpha_{s,k}) - \Psi(\sum_{k''=1}^{K}\alpha_{s,k''}))\right] \\
&- \left[\sum_{k=1}^{K}\sum_{n=1}^{N_s}\pi_{n,k}\log\pi_{n,k}\right],
\end{aligned}
\tag{7}
$$

where $I$ is indication function being either 1 or 0 depending on whether the equation therein is true or not; $\omega_k$ is the $k$-th element in parameter $\omega$, which is set to be symmetric as prior; $\alpha_{s,k}$ is the $k$-th element in $\alpha_s$; $\eta_k^t$, $t \in \{1, 2, ..., T\}$, is the $t$-th element in signature probability $\eta_k$ corresponding to mutation type $t$, and $x_n^t$ is 1 when $x_n$ is observed as mutation $t$, and is 0 if otherwise.

We maximize lower bound with respect to each variational parameters. First, we isolate terms in $L_s(\boldsymbol{\alpha}, \boldsymbol{\pi}; \omega)$ containing $\pi_{n,k}$ and add Lagrange multiplier based on the constraint $1 = \sum_{k=1}^{K}\pi_{n,k}$, then maximize it with respect to $\pi_{n,k}$:

$$
\begin{aligned}
L_s^{[\pi_{n,k}]} &= \left[\sum_{k'=1}^{K} \pi_{n,k'}(\Psi(\alpha_{s,k''}) - \Psi(\sum_{k''=1}^{K} \alpha_{s,k''}))\right] \\
&+ \left[\sum_{k'=1}^{K}\sum_{t=1}^{T} \pi_{n,k'} x_n^t \log \eta_k^t\right] \\
&- \left[\sum_{k'=1}^{K} \pi_{n,k'} \log \pi_{n,k'}\right] \\
&+ \lambda_n(\sum_{k'=1}^{K} \pi_{n,k'} - 1)
\end{aligned}
\tag{8}
$$

Take derivatives with respect to $\pi_{n,k}$, we have:

$$
\begin{aligned}
\frac{\partial L_s^{[\pi_{n,k}]}}{\partial \pi_{n,k}} &= \left[(\Psi(\alpha_{s,k}) - \Psi(\sum_{k''=1}^{K} \alpha_{s,k''}))\right] \\
&+ \left[\sum_{t=1}^{T} x_n^t \log \eta_k^t\right] \\
&- [\log \pi_{n,k} + 1] \\
&+ \lambda_n
\end{aligned}
\tag{9}
$$

Setting this derivative to zero we yield:

$$
\pi_{n,k} \propto \exp\{\Psi(\alpha_{s,k}) - \Psi(\sum_{k''=1}^{K} \alpha_{s,k''}) + \sum_{t=1}^{T} x_n^t \log \eta_k^t\}
\tag{10}
$$

We can further simplify equation(10) since $x_n$ is observed:

$$
\begin{aligned}
\pi_{n,k}^t &\propto \exp\{\Psi(\alpha_{s,k}) - \Psi(\sum_{k''=1}^{K} \alpha_{s,k''}) + \log \eta_k^t\} \\
&= \eta_k^t \exp\{\Psi(\alpha_{s,k}) - \Psi(\sum_{k''=1}^{K} \alpha_{s,k''})\}
\end{aligned}
\tag{11}
$$

Similarly we isolate terms containing $\alpha_{s,k}$ in the lower bound and maximize it with respect to $\alpha_{s,k}$:

$$
\begin{aligned}
L_s^{[\alpha_{s,k}]} &= \left[\sum_{k'=1}^{K} (\omega_{k'} - 1)(\Psi(\alpha_{s,k'}) - \Psi(\sum_{k''=1}^{K} \alpha_{s,k''}))\right] \\
&+ \left[\sum_{k'=1}^{K}\sum_{n=1}^{N_s} \pi_{n,k'}(\Psi(\alpha_{s,k'}) - \Psi(\sum_{k''=1}^{K} \alpha_{s,k''}))\right] \\
&- \left[\log\Gamma(\sum_{k''=1}^{K} \alpha_{s,k''}) - \sum_{k'=1}^{K} \log\Gamma(\alpha_{s,k'}) + \sum_{k'=1}^{K} (\alpha_{s,k'} - 1)(\Psi(\alpha_{s,k'}) - \Psi(\sum_{k''=1}^{K} \alpha_{s,k''}))\right]
\end{aligned}
\tag{12}
$$

Take derivatives with respect to $\alpha_{s,k}$ we yiled:

$$
\begin{aligned}
\frac{\partial L_s^{[\alpha_{s,k}]}}{\partial \alpha_{s,k}} &= \left[(\omega_k - 1)\Psi'(\alpha_{s,k}) - (\sum_{k=1}^{K}\omega_k - K)\Psi'(\sum_{k''=1}^{K}\alpha_{s,k''}))\right] \\
&+ \left[\sum_{n=1}^{N_s}\pi_{n,k}(\Psi'(\alpha_{s,k}) - K\Psi'(\sum_{k''=1}^{K}\alpha_{s,k''}))\right] \\
&- \left[\Psi(\sum_{k''=1}^{K}\alpha_{s,k''}) - \Psi(\alpha_{s,k}) + (\Psi(\alpha_{s,k}) - \Psi(\sum_{k''=1}^{K}\alpha_{s,k''})) + (\alpha_{s,k} - 1)\Psi'(\alpha_{s,k}) - (\sum_{k'=1}^{K}\alpha_{s,k'} - K)\Psi'(\sum_{k''=1}^{K}\alpha_{s,k''})\right] \\
&= (\omega_k + \sum_{n=1}^{N_s}\pi_{n,k} - \alpha_{s,k})\Psi'(\alpha_{s,k}) + \sum_{k'=1}^{K}(-\omega_{k'} - \sum_{n=1}^{N_s}\pi_{n,k'} + \alpha_{s,k'})\Psi'(\sum_{k''=1}^{K}\alpha_{s,k''}) \\
&= (\omega_k + \sum_{n=1}^{N_s}\pi_{n,k} - \alpha_{s,k})\Psi'(\alpha_{s,k}) - \sum_{k'=1}^{K}(\omega_{k'} + \sum_{n=1}^{N_s}\pi_{n,k'} - \alpha_{s,k'})\Psi'(\sum_{k''=1}^{K}\alpha_{s,k''})
\end{aligned} \tag{13}
$$

Setting the equation above to be zero we have one solution as:

$$\alpha_{s,k} = \omega_k + \sum_{n=1}^{N_s}\pi_{n,k} \tag{14}$$

To update posterior parameter $\omega$, we isolate terms containing $\omega$ from the lower bound cross all samples and take derivatives in the following two equations:

$$L^{[\omega_k]} = \sum_{s=1}^{S}\left[\log\Gamma(\sum_{k''=1}^{K}\omega_{k''}) - \sum_{k'=1}^{K}\log\Gamma(\omega_{k'}) + \sum_{k'=1}^{K}(\omega_{k'} - 1)(\Psi(\alpha_{k'}) - \Psi(\sum_{k''=1}^{K}\alpha_{s,k''}))\right] \tag{15}$$

$$\frac{\partial L^{[\omega_k]}}{\partial \omega_k} = \sum_{s=1}^{S}\left[\Psi(\sum_{k''=1}^{K}\omega_{k'}) - \Psi(\omega_k) + (\Psi(\alpha_k) - \Psi(\sum_{k''=1}^{K}\alpha_{s,k''}))\right] \tag{16}$$

To approximate the estimation, we use Newton-Raphson algorithm by using the second derivatives (Hessian matrix) of the lower bound with respect to $\omega_k$. Elements in Hessian matrix is as follows:

$$
\begin{aligned}
\frac{\partial L^{[\omega_k]}}{\partial \omega_k \omega_j} &= \sum_{s=1}^{S}\left[\Psi'(\sum_{k''=1}^{K}\omega_{k''})\right] \\
&= S\Psi'(\sum_{k''=1}^{K}\omega_{k''}), j \neq k, j \in \{1, 2, ..., K\} \\
\frac{\partial L^{[\omega_k]}}{\partial \omega_k \partial \omega_j} &= S\Psi'(\sum_{k''=1}^{K}\omega_{k''}) - S\Psi'(\omega_k), j = k
\end{aligned} \tag{17}
$$

We use $H_\omega$ and $G_\omega$ to denote second derivatives and first derivatives of lower bound with respect to $\omega$ respectively. Using Newton-Raphson algorithm to approximate posterior $\omega$ is in the following:

$$\omega_{new} = \omega_{old} - H_{\omega_{old}}^{-1} G_{\omega_{old}}, \tag{18}$$

Iterate equation(18) until convergence.
